# AgriSeqDB: an online RNA-Seq database for functional studies in agriculturally relevant plant species

**DOI:** 10.1101/330746

**Authors:** Andrew J. Robinson, Muluneh Tamiru, Rachel Salby, Clayton Bolitho, Andrew Williams, Simon Huggard, Eva Fisch, Kathryn Unsworth, James Whelan, Mathew G. Lewsey

## Abstract

**Background:** The genome-wide expression profile of genes in different tissues/cell types and developmental stages is a vital component of many functional genomic studies. Transcriptome data obtained by RNA-sequencing (RNA-Seq) is often deposited in public databases that are made available *via* data portals. Data visualization is one of the first steps in assessment and hypothesis generation. However, these databases do not typically include visualization tools and establishing one is not trivial for users who are not computational experts. This, as well as the various formats in which data is commonly deposited, makes the processes of data access, sharing and utility more difficult. Our goal was to provide a simple and user-friendly repository that meets these needs for datasets from major agricultural crops.

**Description:** AgriSeqDB (https://expression.latrobe.edu.au/agriseqdb), is a database for viewing, analysing and interpreting developmental and tissue/cell-specific transcriptome data from several species, including major agricultural crops such as wheat, rice, maize, barley and tomato. The disparate manner in which public transcriptome data is often warehoused and the challenge of visualizing raw data are both major hurdles to data reuse. The popular eFP browser does an excellent job of presenting transcriptome data in an easily interpretable view, but previous implementation has been mostly on a case-by-case basis. Here we present an integrated visualisation database of transcriptome datasets from six species that did not previously have public-facing visualisations. We combine the eFP browser, for gene-by-gene investigation, with the Degust browser, which enables visualisation of all transcripts across multiple samples. The two visualisation interfaces launch from the same point, enabling users to easily switch between analysis modes. The tools allow users, even those without bioinformatics expertise, to mine into datasets and understand the behaviour of transcripts of interest across samples and time. We have also incorporated an additional graphic download option to simplify incorporation into presentations or publications.

**Conclusion:** Powered by eFP and Degust browsers, AgriSeqDB is a quick and easy-to-use platform for data analysis and visualization in five crops and Arabidopsis. Furthermore, it provides a tool that makes it easy for researchers to share their datasets, promoting research collaborations and dataset reuse.

## Background

RNA-sequencing (RNA-Seq) is currently the preferred technology for genome-wide transcriptional profiling due to its combined ease of use, quality of data and suitability for a diverse range of applications [1, 2]. Recent advances in next generation sequencing (NGS) technologies coupled with decreases in the cost of sequencing have resulted in collection of large volumes of RNA-Seq data from many species [3, 4]. These data are typically deposited in online repositories in formats that are text and/or table-based. Visualization of data is a key early step in transcriptomic analysis for many biologists, allowing examination of data quality, as well as rapid interrogation of leads and hypothesis generation. Many researchers who wish to investigate public transcriptome data are not computational experts, for whom transferring data from the format of online repositories to visualization tools is challenging. This creates a barrier to data reuse. The eFP browser, which was first developed for *in silico* gene expression analysis in Arabidopsis, is an excellent piece of software to display transcriptome data visually [5]. At the time of writing, 20 plant transcriptome datasets are available publicly in dedicated eFP browsers [http://bar.utoronto.ca, 5-17]. Degust is a web-based data visualization tool that provides different functionality from eFP functionality (https://github.com/drpowell/degust). It enables users to view all transcripts form all experiments in an experiment, examine trends between samples, to visualize quality-control metrics and to drill down into subsets of transcripts with expression patterns of interest. These two data browsers could be integrated to provide users with an easy to use tool for accessing and analysing multiple datasets and, by developing some enhanced functionality, they could be used for data download and to generating quality images for presentations or publications.

RNA-Seq is often performed at whole plant or organ level using samples that are composed of different tissues and cell types. This approach masks cell- or tissue-specific information about transcripts, which is important to understand spatial-regulation and functions of genes [18, 19]. Spatial resolution is also important to capture transcripts that are expressed at extremely low levels in specific cell types and that are consequently below the limit of detection in bulk samples of tissues [2]. Temporal gene expression data is also an important tool, which can be used to investigate the mechanisms of genome regulation and to understand the relationships between development and gene function [20]. These approaches have been used in functional studies aimed at deciphering regulatory and structural gene networks of diverse plant species, including forest trees and major crops such as wheat (*Triticum aestivum*), rice (*Oryza sativa*), maize (*Zea mays*), barley (*Hordeum vulgare*), and tomato (*Solanum lycopersicum*) [21-27].

Here we present AgriSeqDB (https://expression.latrobe.edu.au/agriseqdb), a web-based resource that can serve as a public portal for accessing, analysing and visualizing tissue and cell-specific transcriptome dataset from multiple species. Our focus in this implementation is primarily upon transcriptome datasets during the development of seeds and fruits of agriculturally-relevant species. The database integrates two existing open-source browsers and enhances their functionality. The Degust browser provides access to information on genome-wide expression across samples and datasets, aiding the discovery of new genes that can contribute to crop improvement. It also provides quality-control information. The eFP browser allows users to visualize between different samples the abundance of individual transcripts encoded by genes of interest.

## Construction and content

### Database/website architecture

The main structure of AgriSeqDB is described in Fig. 1. It consists of a landing portal that is implemented using an HTML frontend and Python/Django backend to present all datasets and associated meta-data to users. The landing portal allows the user to discover the datasets and navigate to data viewers of interest. The existing eFP browser, which has HTML (frontend) and Python (backend) tools, was selected in order to allow users to view expression data on a gene-by-gene basis [5]. Additionally, the existing tool Degust, is included to allow viewing of expression profiles across all (or a subset of) genes at once [28]. Degust uses an HTML/Javascript frontend and Haskell backend. Both tools were linked and wrapped within the Landing Portal to ensure that users receive a consistent look and feel when using the portal and each viewer (Fig. 2). The source code for the landing portal and integrations with the viewers is available for reuse (https://bitbucket.org/arobinson/agribiohvc). This repository makes use of git submodules to link the source code of eFP and Degust browsers, each of which was modified slightly from original versions to ensure that they link cleanly; source code for modified versions is available at https://bitbucket.org/arobinson/efp and https://github.com/andrewjrobinson/degust, respectively. The Landing Portal and eFP browser use a MySQL database server to store settings and data/meta-data, while Degust uses files on the file system. A central configuration portal was added to ease the loading of data-sets into the database and of the landing portal documentation, allowing organism annotation upload, dataset upload, dataset configuration such as making it private/public, providing external links and abstract etc, and deploying the dataset to what we refer to as GeneView (eFP) or GeneExplore (Degust).

**Figure 1.**
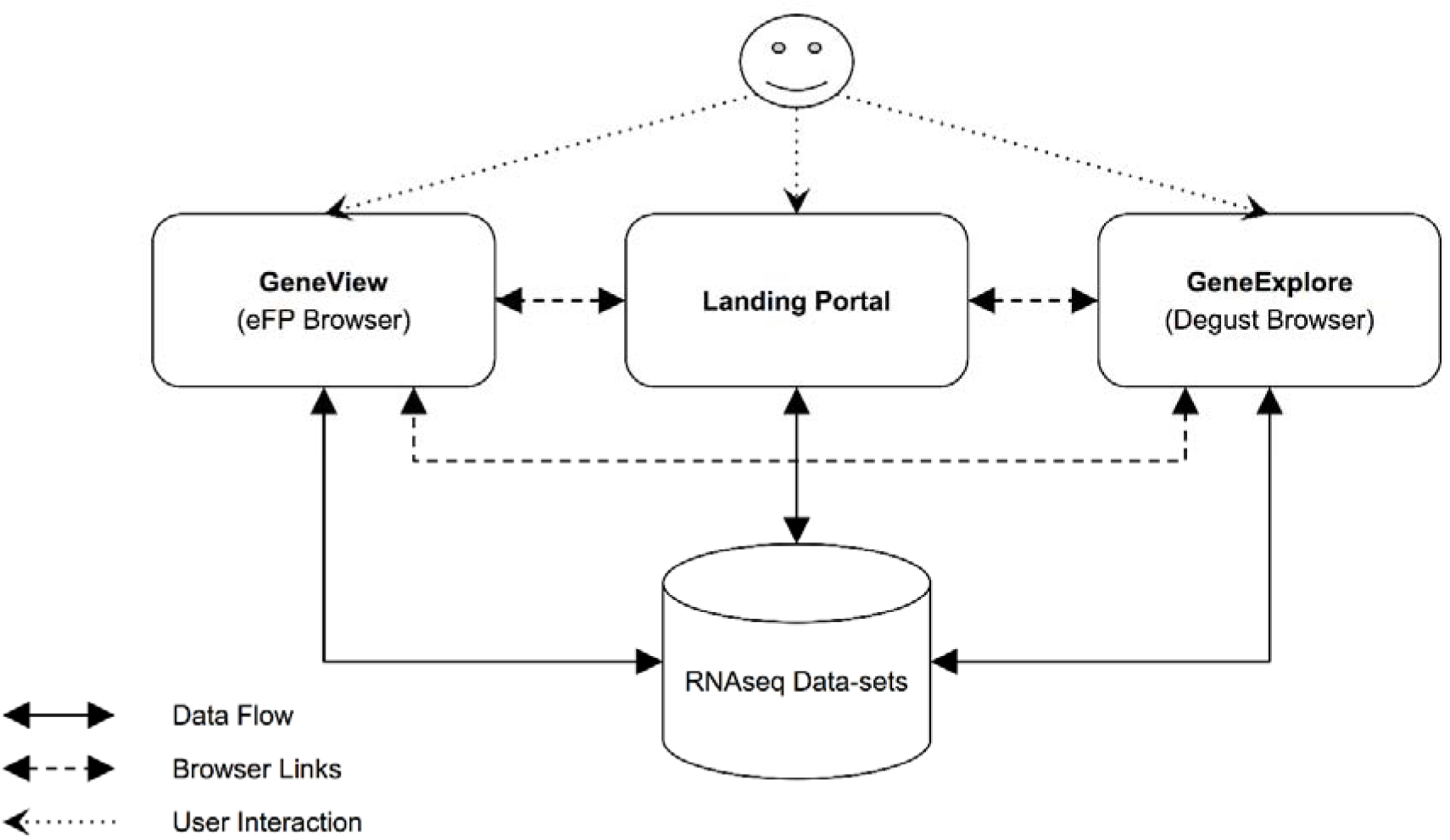
High level structure of AgriSeqDB showing the linkage between data browsers and the central Landing Portal. The Landing Portal provides a central place to access all datasets and provide meta-data that isn’t provided by data browsers. The data browsers provide access to the same data in various forms to enable greater insight.

**Figure 2:**
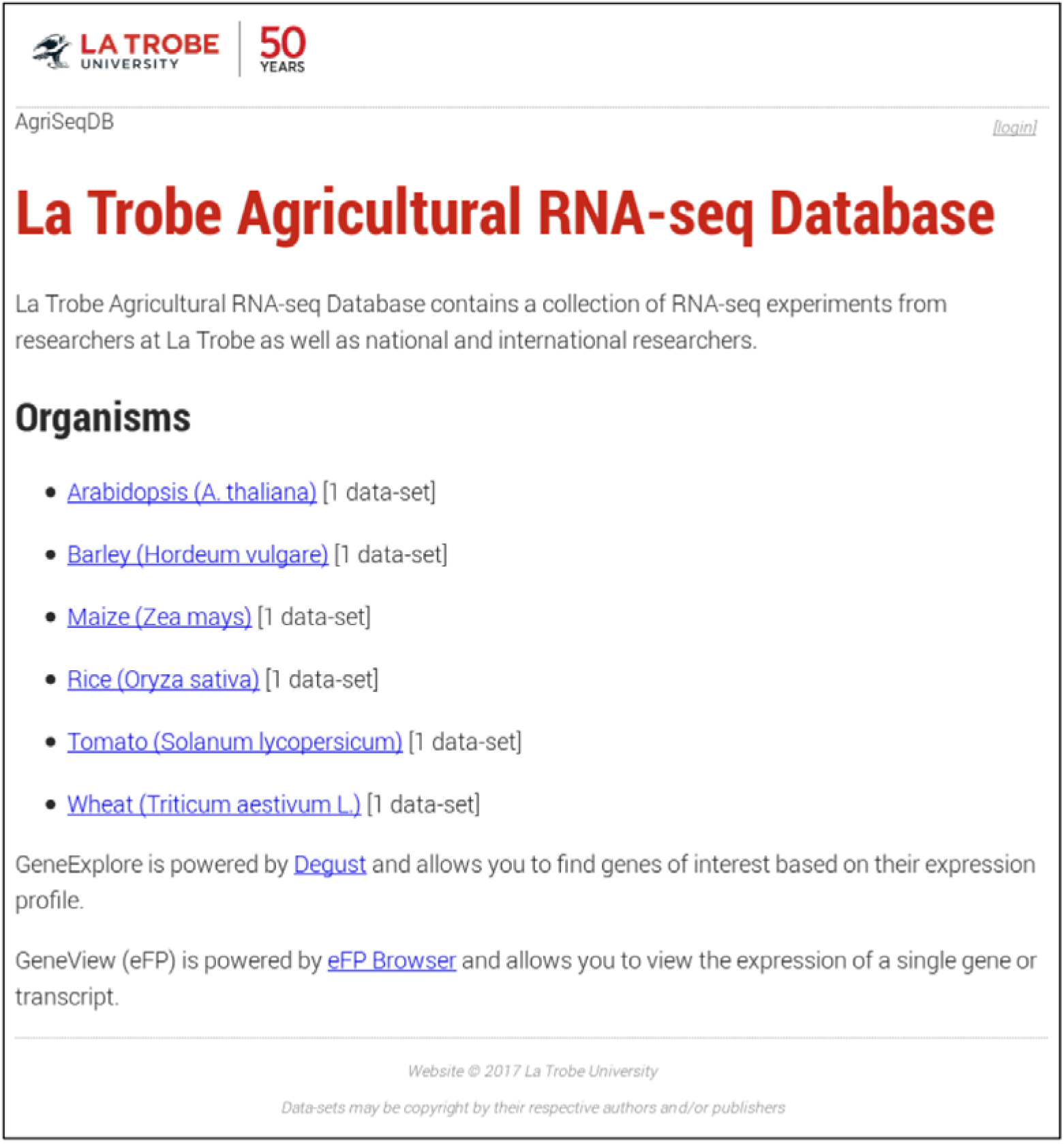
AgriSeqDB home screen showing the six data-sets from species including crops species of major agricultural importance that are currently in the database.

### Data sources

All datasets displayed currently in AgriSeqDB are transcriptomes published recently and deposited in public databases (Fig. 2, Table 1). Users of AgriSeqDB can view data directly from database server without the need to download it and then install/configure a viewer to visualise it. The datasets were generated in six studies of seeds or fruit. The first is a study we conducted of transcriptome changes in whole Arabidopsis seeds during germination, which provides a useful reference due to this species’ high-quality genome sequence and annotation [20]. Additionally, we displayed five datasets from major agricultural crops. These were: A study of transcriptome changes in different tissues (aleurone, starchy endosperm, embryo, scutellum, pericarp, testa, husk and crushed cell layers) of barley grain at different stages of germination [21]; a study on transcriptome changes associated with different cell types of maize endosperm after pollination [22]; a study on transcriptome changes associated with seed germination and coleoptile growth in rice [23]; a study on transcriptome changes associated with fruit development in tomato fruit [26]; and a study on grain/endosperm transcriptome of bread wheat [24].

**Table 1.** RNA-Seq datasets included in AgriSeqDB

| Dataset | Species | Tissue/cell type | Developmental stage/treatment | Data source | Reference |
| --- | --- | --- | --- | --- | --- |
| Seed germination | Arabidopsis<br>(*Arabidopsis thaliana*) | Whole seed | 0 to 48 h post stratification | GSE94459 | [20] |
| Seed germination | Barley<br>(*Hordeum vulgare* L.) | aleurone, starchy endosperm, embryo, scutellum, pericarp testa, husk and crushed cell layers | 0 to 24 h | PRJNA378132 | [21] |
| Endosperm development | Maize<br>(*Zea mays* L.) | Different cell types of endosperm (embryo, nucellus, placento-chalazal region, pericarp, and the vascular region of the pedicel) | 8 d after pollination | GSE62778 | [22] |
| Seed germination and coleoptile growth | Rice<br>(*Oryza sativa* L.) | Embryo and coleoptile | 0 hr to 4 d | (See doi:10.1111/tpj.13418) | [23] |
| Grain/endosperm development | Bread wheat<br>(*Triticum aestivum* L.) | starchy endosperm, aleurone layer, transfer cells | 10, 20, or 30 days post anthesis (DPA) | E-MTAB-2137 | [24] |
| Fruit development | Tomato<br>(*Solanum lycopersicum* L.) | Fruit | Mature ripe fruits | GSE75273 | [26] |

## Utility and discussion

Our goal was to develop a publically accessible transcriptome database that provides simple and readily available tools to perform functional analysis of individual target genes or sets of genes. AgriSeqDB is a highly interactive and multi-view database that can be used for various purposes, including the discovery of genes of interest. Users of AgriSeqDB can view data directly from database server without the need to download it and then install/configure a viewer to visualise it. However, we provide the option for advanced users to download and install their own database for custom datasets.

### GeneView (eFP)

AgriSeqDB also allows users to get a better understanding of individual genes of interest, by inspecting them within GeneView (eFP) (Fig. 3). This incorporates the full existing functionality of eFP [5]. Users can visualise expression of transcripts across all samples so that they may consider the relationships between samples (i.e. growth stage, tissue type, various treatments). Additionally, we incorporated an additional image download function, not previously available. Images may be downloaded in high-resolution.png format for presentations or publications (Fig. 4). This is done by single clicking of the Download button (Fig. 3).

**Figure 3:**
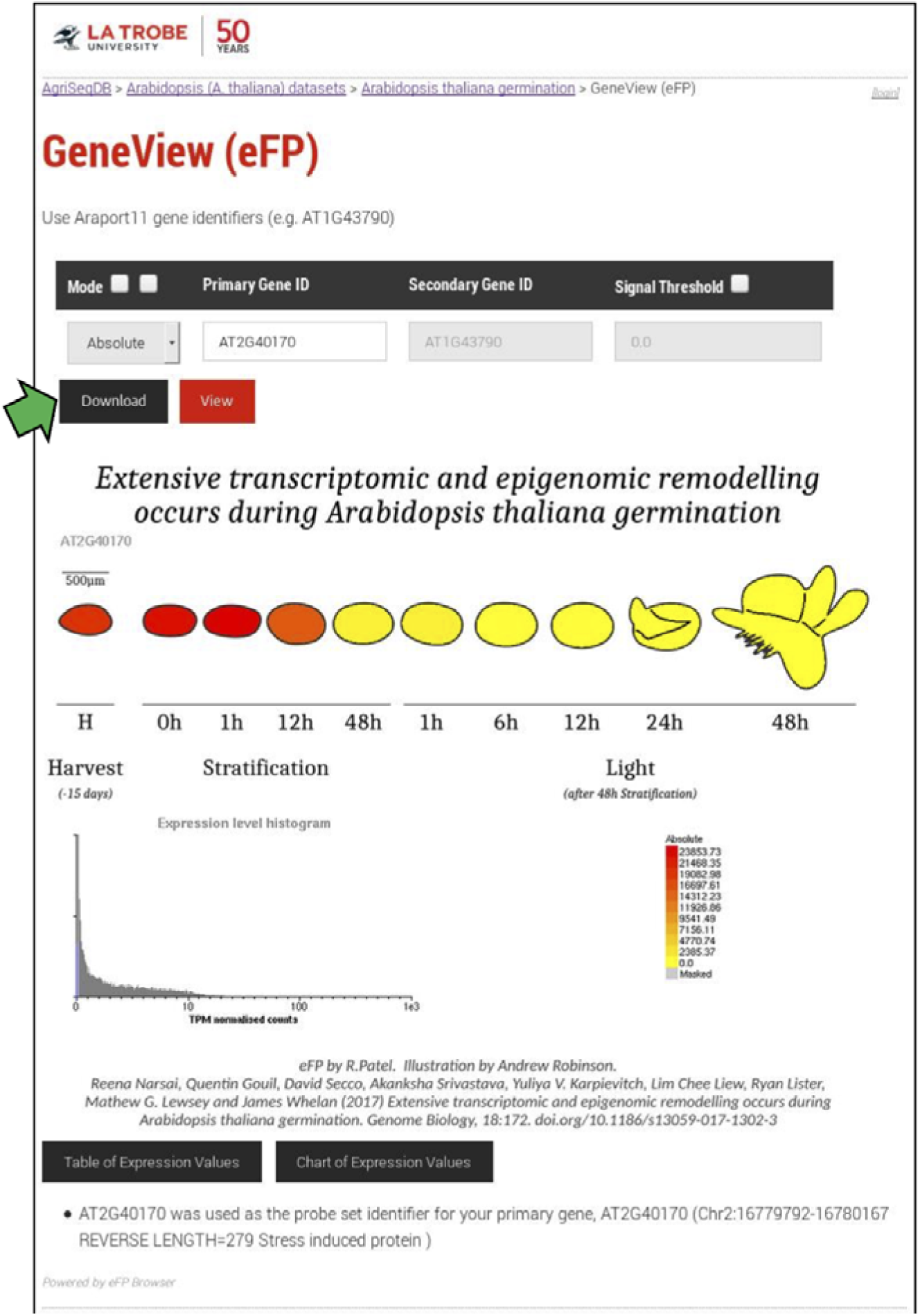
The full screenshot showing AT2G40170 gene expression in GeneView (eFP) browser. The user uses the search form at the top to select the gene of interest and select the mode of operation including: (1) absolute, shows the counts as stored in the database for the primary gene, (2) relative, shows the counts relative to the control for primary gene, and (3) compare, counts as a ratio between the primary and secondary genes. Clicking the view button updates the figure below to show the expression levels of each sample by colour coding the fill area with a scale red-yellow (for absolute) and red-grey-blue (for relative and compare). Alternatively, the user can click the download button (indicated by a green arrowhead) to download the expression image at twice the resolution as shown on-screen (ready for publication) as shown in Fig. 4. Data is from transcriptome of Arabidopsis seeds during germination [20].

**Figure 4:**
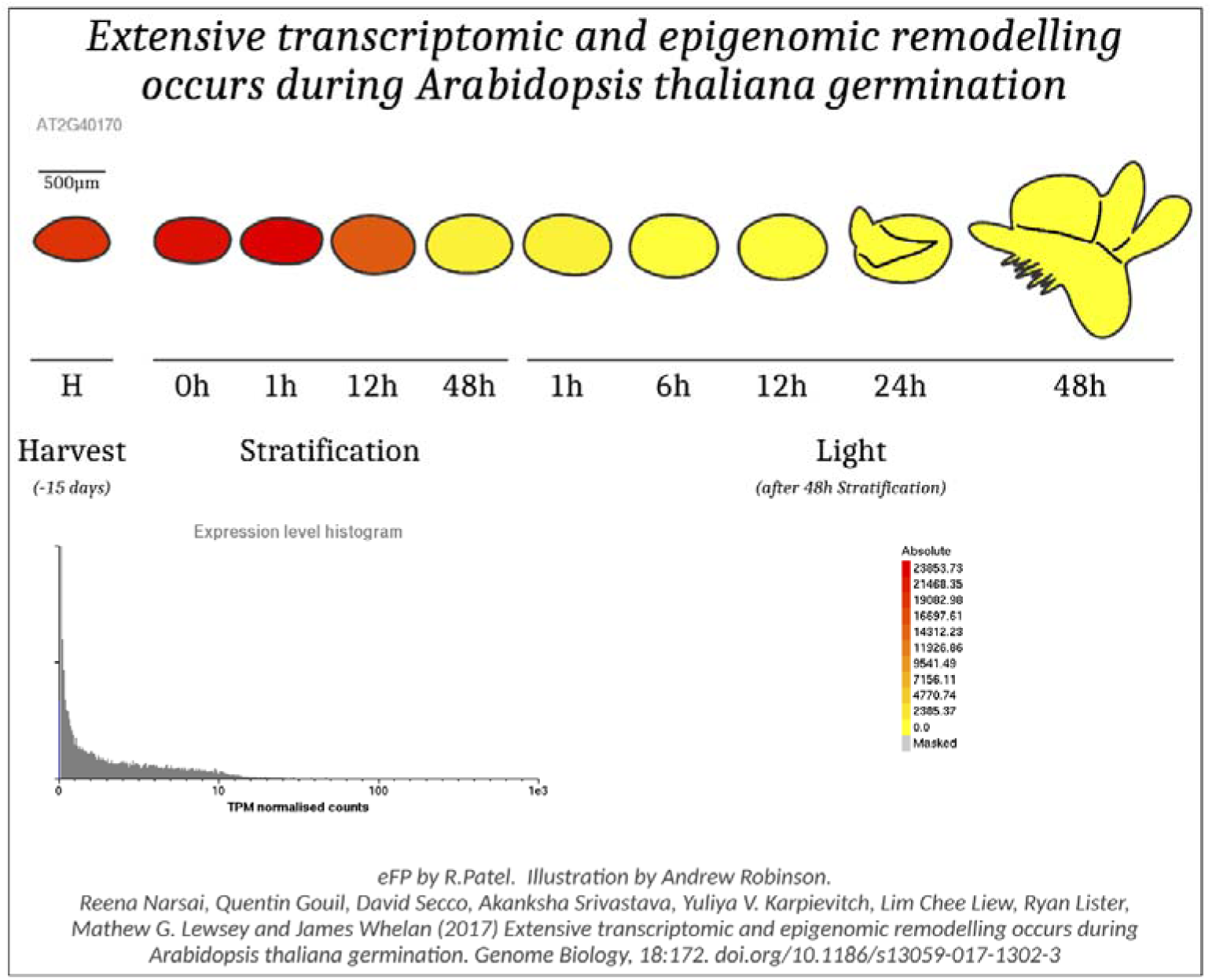
A high-quality image for AT2G40170 gene expression downloaded in GeneView (eFP) Browser. Data is from transcriptome of Arabidopsis seeds during germination [20].

### GeneExplore (Degust)

Users are presented with a simple interface to query all genes using GeneExplore (Degust) (Fig. 5a). Extensive existing functionality is available to users within Degust [28]. Filters can be created on the data based upon expression levels in individual samples, false discovery rate (FDR) and Log2 fold-change cut-off. Sub-sets of samples or transcripts can be selected for analysis can be analysed and the sample for referencing fold-change can be selected. MA plots of comparisons between pairs of samples can be displayed (Fig. 5b). Data quality metrics can be assessed by inspecting whether the replicates of each sample group together in the multidimensional scaling (MDS) plot (Fig. 5c). Data tables can also be downloaded for selected transcripts in.csv format for downstream analyses.

**Figure 5:**
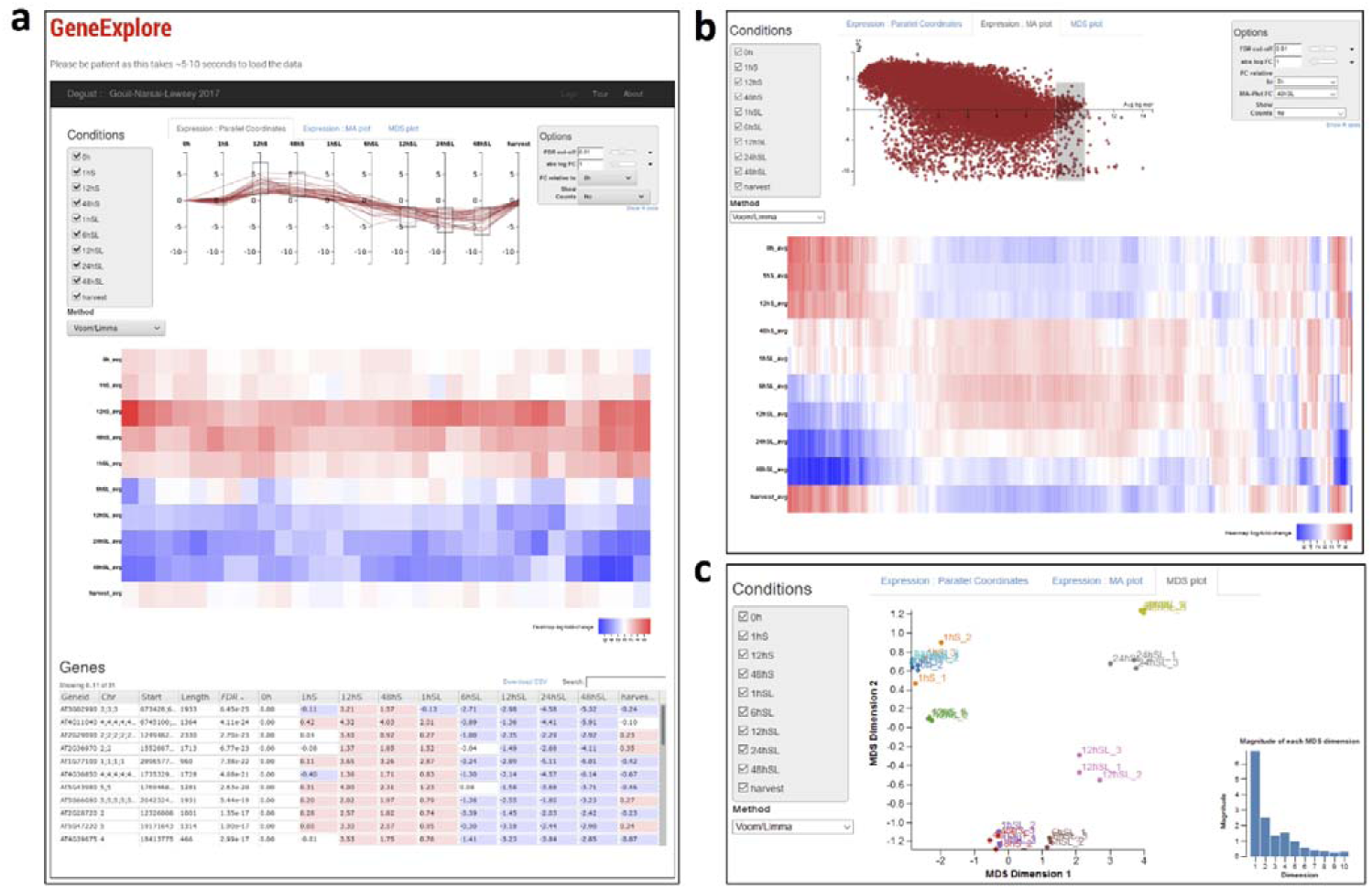
Screen shot of the GeneExplore (Degust) browser and subsequent result pages. (**a)** the Arabidopsis time-series dataset is shown here as an example, displaying transcripts that are up-regulated in S samples but down-regulated in SL samples (Top panel). The user can select which samples they wish to see with the checkboxes in the top left of screen along with the method of analysis (voom/limma, edgeR, or voom). In the top right, the user can control the rendering and thresholds of using the options dialog. All genes that match filters above are shown in a heat-map, which clusters genes with similar levels of expression (Middle panel). Running the mouse-over each gene highlights it in the plots above. Table showing all matching genes in tabular format with the expression levels for each sample, false discovery rate and any extra annotation columns provided in the dataset (Lower panel). In the top centre the user can limit genes by using 1 of 3 interactive plots, and the parallel coordinates plot allows the user to limit genes by their log fold gene expression (per sample). (**b)** Example of an MA plot. Users can limit genes by drawing a box around genes on the on the MA plot; the two samples used for the MA plot are specified in the Options dialog (top right). (**c)** An MDS plot showing groupings of the individual replicates of each sample. Data is from transcriptome of Arabidopsis seed during germination [20.

### Data-set Administration (Advanced Usage Case)

One key inclusion to AgriSeqDB is the data-set administration tool, available to advanced users upon download and install of their own AgriSeqDB. eFP browser did not contain an interface to upload data, so configuration required much manual interaction. While Degust contained its own administration tool, it was not flexible enough to accommodate eFP and the landing portal. Consequently, we developed a new data upload interface to encompass both eFP and Degust. This allows the user (secured by username/password) to upload new datasets and deploy them to each of the viewers. An example configuration is included in Supplementary Figure 1.

## Conclusions

We believe AgriSeqDB will be an important resource and data-reuse tool for plant biologists who seek greater insights into the role of individual genes or group of genes in biological processes, including for comparative studies in crop species of major agricultural importance. The databases will be periodically updated with more viewers and datasets, focusing on additional tissue and cell-specific datasets from crop species. The database currently contains results of RNA-Seq from different tissues and cell types, and it is planned that transcriptome data from single cell RNA-Seq will be added in the future. In the long term it is envisaged that users will be provided with links to GEO auto-download and view as well as allowed to upload datasets at least temporarily. All source code is freely available for reuse by advanced users.

## List of abbreviations

eFP: Electronic Fluorescent Pictograph
FDR: False Discovery Rate
GEO: Gene Expression Omnibus
HTML: Hypertext Markup Language
MDS: Multi-Dimensional Scaling
NGS: Next Generation Sequencing
RNA-Seq: RNA Sequencing

## Acknowledgments

We gratefully acknowledge the excellent work of Dr Nicholas Provart (U. Toronto) and the eFP team past and present as well as the developer of Degust, David Powell (Monash University), whose software we utilised and extended here. We also thank them for their advice when establishing our database. We thank all members of the teams who generated the data we have displayed in AgriSeqDB, who are too numerous to list here. We thank Maoshan Chen (La Trobe University) for his help with data processing.

## Availability of data and materials

The database is freely available via https://expression.latrobe.edu.au/agriseqdb. It is compatible with all modern popular web browsers and possible to use by tablets and mobile/cell phones. Database source code is available for reuse at https://bitbucket.org/arobinson/agribiohvc. Modified Degust and eFP source code used in this project is available at https://bitbucket.org/arobinson/efp and https://github.com/andrewjrobinson/degust.

## Authors’ contributions

AJR developed the data portal, created illustrations (for eFP) and uploaded data. MT contributed to project development. MGL and JW conceived the project and provided scientific direction. MGL conducted tool research. CB annotated Meta-data and RDA records. RS, AW, SH and EF participated in project management. AJR, MT, MGL and JW wrote the manuscript. All authors read and approved the final manuscript.

